# Neuroendocrine Stress Induces Differential Oxidative Stress and Antioxidant Profiles between Proactive and Reactive Stress Coping Styles

**DOI:** 10.64898/2026.02.03.703382

**Authors:** Princess Sunday-Jimmy, Robert J. Fialkowski, Brady J. Bush, Peter D. Dijkstra, Ryan Y. Wong

**Affiliations:** University of Nebraska at Omaha Biology Department, Omaha, NE USA; Central Michigan University Biology Department, Mount Pleasant, MI USA

**Keywords:** neuroendocrine stress, oxidative stress, stress coping styles, antioxidants

## Abstract

Neuroendocrine stressors can disrupt the brain’s redox equilibrium by generating high levels of reactive oxygen species (ROS) that lead to oxidative stress. The magnitude of the effect of neuroendocrine stressors on brain redox equilibrium can be influenced by many internal and external factors. To what extent the relationship between neuroendocrine and oxidative stress is modulated by an individual’s stress coping style is only beginning to be understood. To explore this, we subjected proactive and reactive zebrafish to an acute novelty stressor and subsequently quantified changes in behavior and whole brain biomarkers of oxidative stress and antioxidants (DNA damage, total glutathione (GSH), glutathione ratio, oxygen radical absorbance capacity (ORAC), and superoxide dismutase (SOD). Stressed fish had significantly higher total glutathione, trends higher ORAC, DNA damage, and glutathione ratio, and trend for lower SOD levels compared to controls. In addition, individuals with a reactive stress coping style exhibited significantly higher levels of SOD and glutathione ratio, and a trend for ORAC compared to proactive individuals. From a principal component analysis, we also found that the reactive individuals had significantly higher PC1 scores (antioxidant axis) compared to the proactive, and a trend for stressed fish having higher PC1 scores than control. The oxidative stress axis (PC2) showed that the stressed fish had a significantly higher PC2 score relative to control fish. Our results show that neuroendocrine stress-induced disruption of redox equilibrium in the brain differs by stress coping style. Those with a reactive stress coping style have elevated antioxidant capabilities and capacities. Overall, our findings suggest that elevated reactivity to neuroendocrine stressors commonly seen in reactive stress coping styles may be mitigated through the glutathione buffering system and other antioxidants.

## Introduction

Animals face a variety of stressors throughout their lives, and their behavioral and physiological responses to overcome these stressors differ among individuals [1,2]. Stress-induced increases in glucocorticoids and catecholamines can lead to oxidative stress [3]. These hormones increase metabolic activity and excitatory signaling, which leads to higher mitochondrial production of reactive oxygen species (ROS) and could inhibit antioxidant responses [4–8]. Oxidative stress arises from an imbalance between the production of reactive oxygen species (ROS) and the antioxidant defense system, disrupting redox homeostasis and promoting damaging effects on the structural and functional aspects of the cell [9]. The degree of disruption from neuroendocrine stress exposure depends on individual characteristics, including their stress-coping mechanisms [10].

Stress-coping styles are characterized by correlated suites of behaviors and physiological responses to a neuroendocrine stressor that are consistent across time and contexts [11–14]. For the proactive-reactive stress coping spectrum, those with a proactive stress coping style tend to be more aggressive, bold, active, and with lower neophobia. Physiologically, proactive coping styles are characterized by lower cortisol levels and neuroendocrine stress reactivity, and higher sympathetic reactivity. The reactive stress coping styles exhibit the direct opposite characteristics [11,15].

Despite neuroendocrine stress axis activity and reactivity directly altering oxidative stress and differing by stress coping style, the link between stress coping styles and oxidative balance is not well characterized [16]. A study using selectively bred mouse lines that differed in attack latency showed higher levels of serum antioxidant capacity but not reactive oxygen metabolites in the long attack latency line (presumably a reactive stress coping style) [17–19]. In contrast, highly neophobic birds (i.e., reactive stress coping style) had lower plasma antioxidant capacity and higher oxidative stress than those with lower neophobia [20]. Altogether, oxidative stress and antioxidant activity may vary between stress coping styles, but this effect is complex and tissue-specific, and highly context-dependent (e.g., whether the animal experiences stress or not)[17,21–23]. Studies to date investigating the relationship between stress coping styles and oxidative profiles have mainly used peripheral components such as blood and liver. However, the oxidative profiles between blood and brain are not consistent [24–26]. Despite the brain being key to coordinating behavioral and physiological processes in a stress coping style, a target of glucocorticoids, and highly vulnerable to oxidative stress, little is known in the brain about neuroendocrine-induced oxidative stress between stress coping [27,28].

Therefore, this study aims to investigate how an individual’s stress coping style can influence disruption of brain redox homeostasis under acute neuroendocrine stress. We hypothesize that there would be differences in brain oxidative stress and antioxidant responses between the two coping styles to acute neuroendocrine stress. We predict that individuals with the reactive stress coping style would exhibit increased levels of markers for oxidative stress. To test this prediction, we compared brain oxidative profiles of zebrafish selectively bred to display the proactive and reactive stress coping styles in both stressed and baseline conditions.

## Methods

### Subjects

We used zebrafish from lines selectively bred to display, on average, the proactive and reactive stress coping style [14,29]. We randomly chose 24 zebrafish (12 baseline and 12 stressed fish) for each line that were selectively bred for 13 generations and were 4-8 months old at the time of testing. Some samples were used for oxidative stress and antioxidant assay optimizations and were not included in the analysis. Final sample sizes consisted of 9 baseline reactive (4 females, 5 males), 10 stressed reactive (3 females, 7 males), 12 baseline proactive (8 females, 4 males), and 10 stressed proactive (6 females, 4 males) individuals. Sex was determined upon visualization of the testes or ovaries on dissection. Prior to testing, all fish were housed in mixed-sex 40-litre tanks on a custom-built recirculating system with solid filtration. Fish were maintained under a 14:10 light/dark cycle, temperature (26-27 °C), water quality (pH: 6.8-7.4, conductivity: 600-900 μS), and fed twice daily with Tetramin Tropical Flakes (Tetra, USA). All procedures were approved by the University of Nebraska Omaha Institutional Animal Care and Use Committee (17-070-09-FC).

### Acute Neuroendocrine Stressor

We induced an acute neuroendocrine stress response to a novelty stressor in our stressed proactive and reactive groups by placing fish individually in a novel open field environment for 30 minutes. We quantified stress behaviors in this open field test (OFT) following established protocols [1,30]. In brief, each fish was placed in a 31.75cm x 31.75cm acrylic tank with black walls and a white bottom, and filled with 4 liters of water. We video recorded the trial for 30 minutes and later quantified behaviors using commercial software (Ethovision Noldus XT Version 17). To be consistent with other studies using OFT to measure response to a novelty we only analyzed the first five minutes of the video recordings [1,31–34]. We quantified time frozen (sec), total distance swam (cm), and average swim velocity (cm/sec). Freezing behavior was defined as the duration of time a fish moved slower than 0.5 cm/sec. After 30 minutes in the open field test, we immediately extracted and froze the brain of each fish on dry ice and stored it at -80°C until tissue processing. Non-stressed (baseline) fish were caught directly from their home tank and had their brains extracted and stored in the same manner.

### Choice of markers of oxidative stress

In each brain sample, we measured various markers of oxidative stress and antioxidant capacity. We assessed total antioxidant capacity (TAC) as a measure of both enzymatic and non-enzymatic antioxidant capacity. We also measured glutathione since it is one of the most abundant small molecule antioxidants. We used an assay measuring both reduced and oxidized glutathione levels which also provides us with an assessment of oxidative stress levels. We evaluated oxidative DNA damage by measuring levels of 8-hydroxy-2-deoxyguanosine (8-OhDG). Below is a description of all the assays. For each assay, we ran samples in duplicate and most assays included inter-assay controls. All samples were processed by researchers that were blinded with respect to treatment. For all assays we used clear flat-bottom 96-well plates unless indicated otherwise.

### Whole brain quantification of oxidative stress and antioxidant levels

#### Tissue preparation

We homogenized brain sample and prepared different aliquots of brain homogenate for total antioxidant capacity (TAC), superoxide dismutase (SOD), glutathione, and oxidative DNA damage assay. Brains were homogenized in a homogenization buffer (50 mM potassium phosphate buffer, 0.5mM EDTA, pH 7.0) plus protease inhibitor cocktail (Sigma-Aldrich P2714, dissolved in 10 mL water, diluted 1:10 in homogenization buffer) and divided into separate aliquots for further analyses using previously described protocols [25,35]. Homogenization was carried using a Cole-Parmer Battery-Operated Pestle Motor Mixer (Cole-Parmer North America, Vernon Hills, Illinois) at a concentration of 10 µL homogenization solution per 1 mg tissue. Once homogenized, the brain sample homogenates were split into three separate aliquots (40 µL, 60 µL, 100 µL).

Aliquots of brain homogenate were created for each assay as follows: The 100 µL homogenates were centrifuged at 4°C and 14,000g for 10 minutes. Following centrifugation 60 µL of supernatant was removed and stored at -80°C until SOD analysis, while the remaining supernatant was discarded and the pellet was stored for quantifying DNA damage. The 40 µL aliquot was diluted 1:1 with lysis buffer (20 mM Tris–HCl, 137 mM NaCl, 1% NP-40, 10% glycerol, 2 mM EDTA) and centrifuged at 4°C and 10,000g for 10 minutes. After centrifugation, 65 µL of the supernatant was aliquoted into a 0.5 mL centrifuge tube. From this aliquot, 14 µL was transferred to a new tube for usage in a bicinchoninic acid (BCA) assay. Both aliquots were stored at -80°C until analysis of oxygen radical absorbance capacity (ORAC) to assess TAC. The 60 µL aliquot was centrifuged at 4°C and 10,000g for 10 minutes. Once centrifuged, 45 µL of supernatant was aliquoted into another 0.5mL tube and diluted 1:1 with 5% 5-sulfo-salicylic acid dihydrate (SSA) solution (Sigma-Aldrich 247006) which was again centrifuged at 4°C and 10,000g for 10 minutes. Upon centrifugation, 70 µL of the supernatant was aliquoted into a 0.5 mL centrifuge tube. From this aliquot, 14 µL was transferred to a new tube for a BCA assay. Both aliquots of supernatant were stored at -80°C until analysis of glutathione.

#### Protein quantification

Protein concentration of each prepared brain homogenate (excluding DNA damage) was measured with a bicinchoninic acid (BCA) Protein Assay kit (Pierce, Rockford IL) following the manufacturer’s protocol. Frozen supernatant was thawed on ice and diluted 1:4 with buffer used in their respective sample preparation. We used 10 µL of this diluted supernatant in the assay. Absorbance was read by a plate reader (Epoch2T, Biotech Instruments, Winooski, VT, USA).

#### Superoxide dismutase (SOD)

Superoxide Dismutase (SOD), an enzymatic antioxidant, was measured via a modified competitive assay (Dojindo Molecular Technologies, Inc.) as previously described [35]. We used 20 µL of prepared brain sample homogenate in duplicate in a 96-well plate at a range of different concentrations and mixed with 200 µL WST-1 working solution and xanthine oxidase. We used 4 or 5 different concentrations of prepared brain sample homogenate, ranging in protein concentration from 0.03 – 0.12 ng/ µL. The reaction rate was read over five minutes and analyzed for the IC50, or where 50% of the reaction is inhibited by SOD at a set concentration of sample and reported per unit of SOD (one unit of SOD is defined as the amount of enzyme in 20 µL of sample solution that inhibits the reduction reaction of WST-1 with superoxide anion by 50%). Absorbance was read by a plate reader (Epoch2T, Biotech Instruments, Winooski, VT, USA).

#### Total antioxidant capacity

Total antioxidant capacity (TAC) was measured in 20 µL of prepared brain sample homogenates via an Oxygen Radical Absorbance Capacity (ORAC) assay as previously described [36]. We standardized the samples to a protein concentration of 150 µg/mL. TAC is reported as µmol Trolox equivalent (TE)/µg protein. Absorbance was read by a plate reader (Spectramax M3, Molecular Devices, Sunnyvale, CA, USA). The intra-assay and inter-assay coefficient of variation was 1.4% and 4.6%, respectively.

#### Glutathione

We measured glutathione using a glutathione fluorescence kit (Arbor Assays, Ann Arbor, MI, USA) per manufacturer’s protocol. Samples were first diluted 1:2.5 with kit assay buffer to reduce SSA concentration to 1%. Fifty microliters of treated samples, standards, and blanks were plated in duplicate in a black half-area 96-well plate, and 25 μL of Thiostar detection reagent was added to each well. Plates were incubated at room temperature for 15 min before endpoint fluorescent emission at 510 nm with excitation at 390 nm was read by a plate reader (Tecan M200 Infinite) to quantify the concentration of free reduced glutathione (fGSH). Next, the concentration of oxidized glutathione dimers (GSSG) was quantified by adding 25 μL kit reaction mixture to each well followed by a second 15-min incubation to convert all GSSG in the sample to fGSH before a second fluorescence reading to find the total concentration of reduced glutathione (tGSH) in each sample. GSSG was calculated as (tGSH— fGSH)/2. We used the ratio of fGSH:GSSG as an overall marker of oxidative stress with higher ratios corresponding to less oxidized glutathione and hence lower levels of oxidative stress. All samples are reported standardized to protein concentration in the sample (total GSH / protein concentration). The intra-assay coefficient of variation was 2.4% (samples were measured using one plate).

#### Oxidative DNA damage

We measured oxidative DNA damage (8-OHdG) following a previously described protocol with modifications to allow for measurements of small tissue samples [35]. DNA was extracted from homogenized brain tissue samples using a DNA extraction kit (Zymo quick-DNA miniprep plus kit, Irvine, CA, USA) per manufacturer’s protocol. Purified DNA samples were extracted from the columns by adding 100 µL of DNA elution buffer and incubating at room temperature for 5 minutes, followed by centrifugation at 17,000 × g for 1 minute. This step was then repeated with an additional 100 µL of DNA elution buffer, again incubating for 5 minutes at room temperature and centrifuged at 17,000 × g for 1 minute. This two-step elution procedure was used to increase the amount of extracted DNA. We then used ethanol precipitation to increase DNA concentration in each DNA sample through addition of 19 µL 3M sodium acetate (pH 5.2) and 475 µL cold ethanol. Samples were then stored at -20 °C overnight before undergoing 30-minute centrifugation at top speed at 4 °C. All ethanol was then removed, and the DNA was reconstituted in 15 µL elution buffer. After ethanol precipitation, the total DNA concentrations were determined using a BioTek take3 reader in an Epoch2T microplate reader using the Nucleic Acid Quantification software provided. We standardized all DNA samples to 100 ng/µL and stored sample at 4 °C until digestion.

DNA samples were digested following a modified protocol from Quinlivan and Gregory [37]. Extracted DNA were added to a digest mix [173.61 Units benzonase, 208.33 mUnits phosphodiesterase, and 138.89 Units phosphatase in a tris-HCl buffer (20 mM, pH 7.9) containing 100 mM NaCl and 20 mM MgCl_2_] at a concentration of 4 µL digest mix per 0.8 ng DNA and incubated at 37 °C overnight to ensure complete hydrolysis of DNA samples. Following digestion, samples were stored at -20 °C until use in DNA damage assay.

DNA damage (8-OHdG) was measured with a DNA damage ELISA kit (StressMarq, Biosciences Inc., Victoria, BC, Canada) following manufacturer’s protocol. Results are reported as ng damage (8-OHdG) / ng DNA.

### Statistical Analysis

All statistical analyses were performed in SPSS (Version 24). We used a general linear model to assess for the main effects of treatment, stress coping styles, and sex, and treatment by stress coping style by sex interaction effect on each oxidative stress and antioxidant marker (SOD, ORAC, total GSH, glutathione ratio, and DNA damage). We only found sex to have a significant main effect on ORAC levels and reported results with sex both included and excluded. In all other variables, there was no significant main effect of sex or treatment by stress coping style by sex interaction effect (Supplementary Table 1). Thus, we removed sex from the model and report the main effects of treatment, stress coping style, and treatment by stress coping style interaction effect. We also conducted a principal component analysis (PCA) on the five oxidative stress and antioxidant markers to obtain principal component scores for evaluation in a general linear model as described above. As we only had behavioral measures for stressed fish, we used a general linear model to assess for main effects of stress coping styles and sex, and a stress coping style by sex interaction effect on total distance swam, average swim velocity and time frozen. There was no significant main effect of sex or stress coping style by sex interaction effect (Supplementary Table 2). Thus, we removed sex from the model and report the main effect of stress coping style on each behavior. To investigate relationships between individual variation of behavior and brain oxidative stress and antioxidant quantities, we ran linear mixed models to examine for effects of redox biomarker, stress coping style, and redox biomarker by stress coping style interaction effect.

## Results

### Effect of Acute Stress and Coping Styles on Oxidative Stress and Antioxidant Markers

#### Oxidative Stress Markers (DNA damage & Glutathione Ratio)

We found that stressed fish had a trend for higher DNA damage than control fish (stress treatment effect: *F*_l,35_=3.560, *p* = 0.068 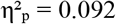), [(Fig. 1a). This effect of stress treatment on brain DNA damage did not vary by stress coping style (strain by stress treatment interaction:*F*_l,35_ = 0.447, *p* = 0.508,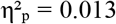). DNA damage levels were not different between stress coping styles (strain effect: *F*_l,35_ =1.798, *p* = 0.189, 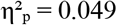). For the glutathione ratio, we found that stressed fish tended to have a higher glutathione ratio level than control fish (stress treatment effect: *F*_l,36_ = 3.291, *p* = 0.078, 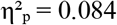) (Fig. 1b). This effect of stress treatment on brain glutathione ratio did not differ by stress coping style (strain by stress treatment interaction: *F*_l,36_ = 0.150, *p* = 0.700, 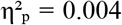). Glutathione levels did not vary between stress coping styles (strain effect: *F*_l,36_ = 0.869, *p* = 0.358, 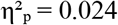).

**Figure 1.**
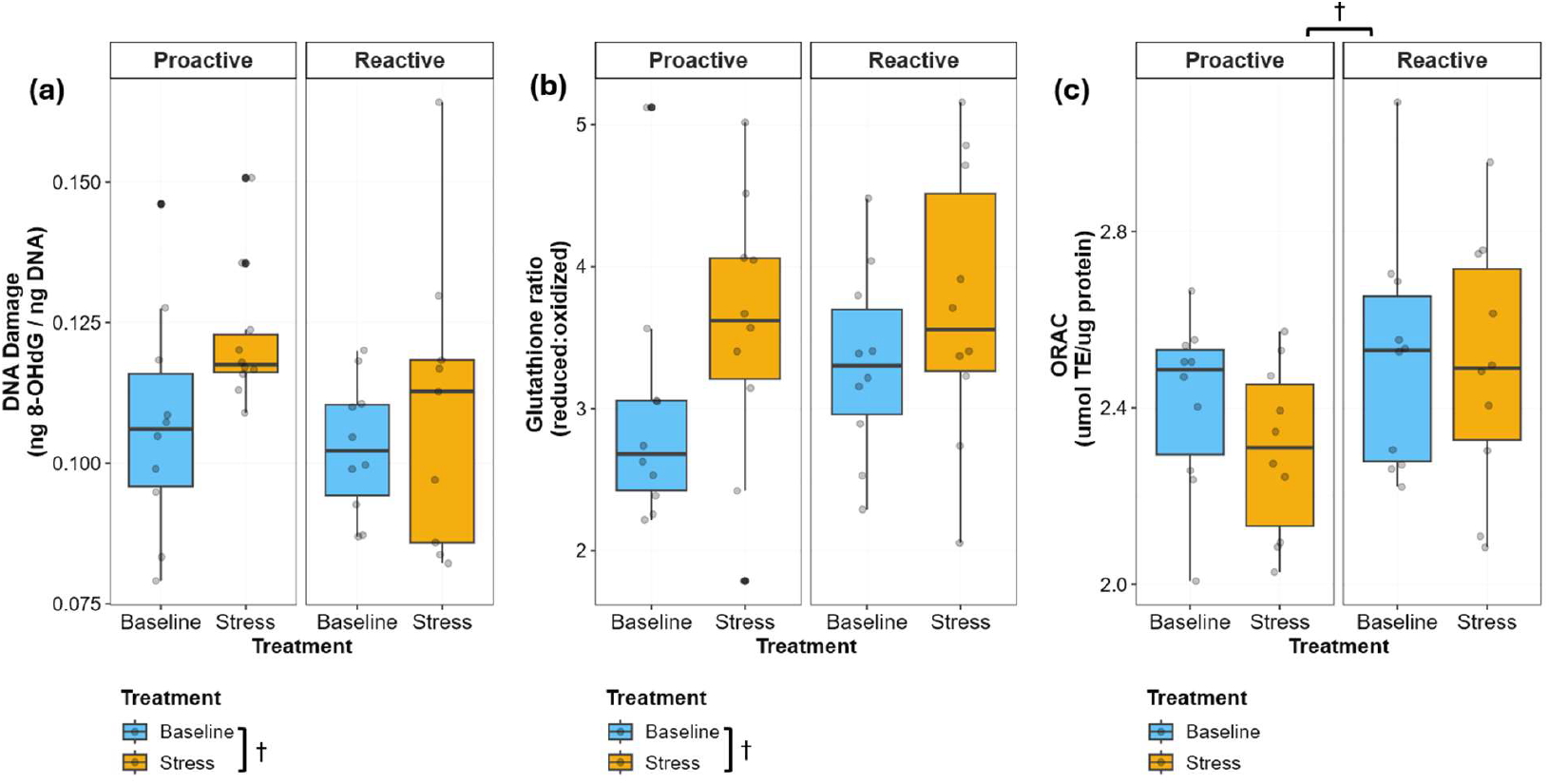
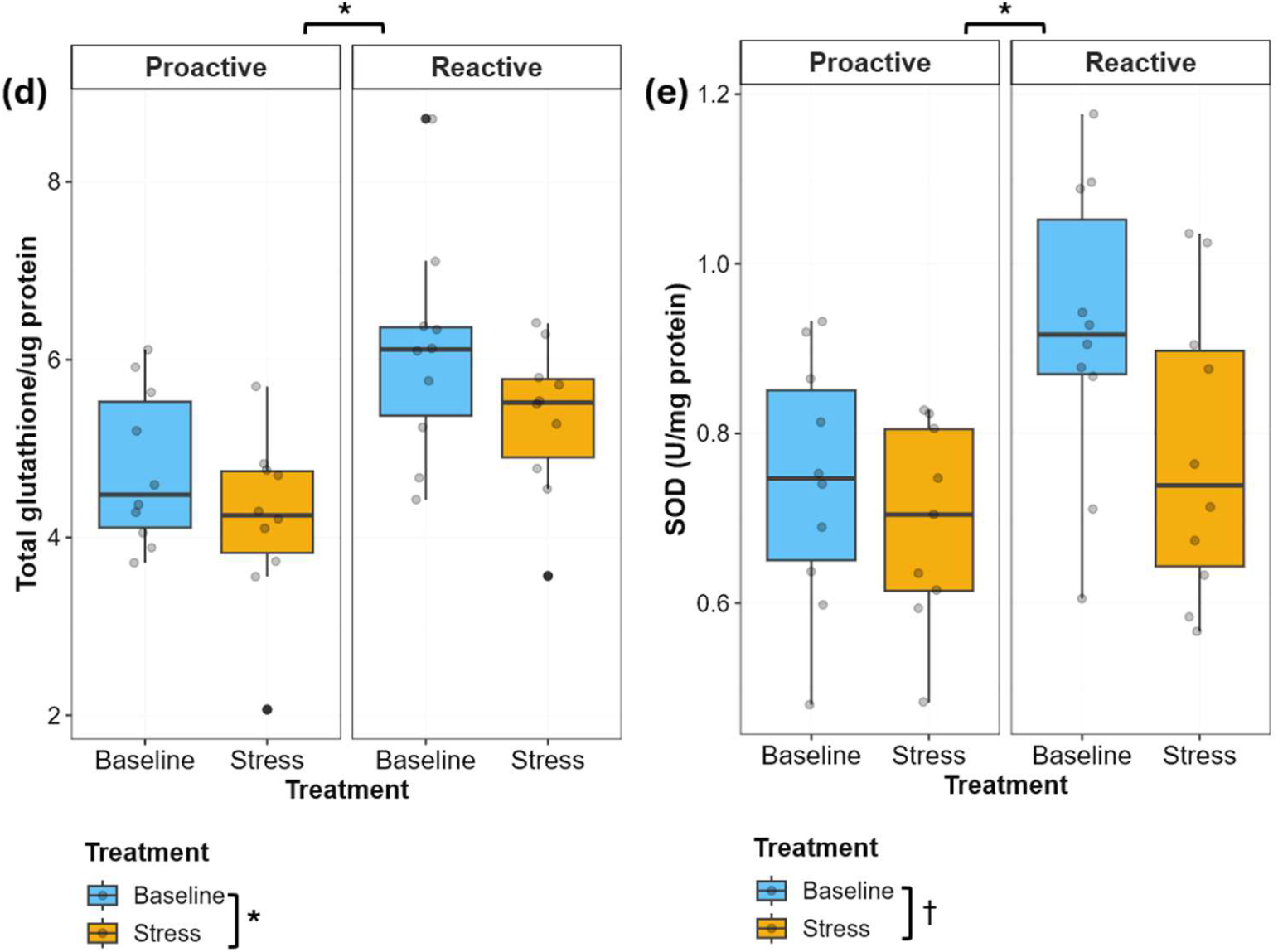
Effects of acute stress on oxidative stress and antioxidant markers across stress coping styles. The boxplot shows (a) DNA damage, (b) glutathione ratio (c) ORAC, (d) total glutathione, and (e) SOD levels. The blue box indicates the baseline group of fish, while the yellow box represents the stressed group of fish. (*) represents *p* < 0.05 and (†) represents 0.05 < *p* < 0.10. Each dot represents a fish.

### Antioxidant Markers (ORAC, SOD & Total Glutathione)

We found that females had significantly higher levels of ORAC than males (sex effect: *F*_l,32_ =9.384, *p* = 0.004,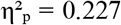). ORAC levels were significantly higher in the reactive relative to the proactive (strain effect: *F*_l,32_ =6.742, *p* = 0.014, 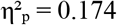). With sex removed from the model, we observed that the reactive line had a trend for higher levels of ORAC than the proactive line (strain effect: *F*_l,36_= 3.766, *p* = 0.060, 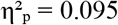) (Fig. 1c). This effect of stress coping styles on the brain ORAC did not differ by treatment levels (strain by stress treatment interaction: *F*_l,36_ = 0.356, *p* = 0.555, 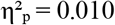). ORAC levels did not vary between treatment levels (stress treatment effect: *F*_l,36_ =0.746, *p* = 0.394, 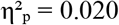). For total GSH, we observed that the stressed fish had significantly higher total glutathione levels (Fig. 1d) relative to control fish (stress treatment effect: *F*_l,36_ =4.453, *p* = 0.042, 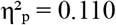). We found that the reactive fish had significantly higher total glutathione levels compared to the proactive fish (strain effect: *F*_l,36_=15.261, *p* = 0.001, 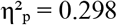). This effect of stress coping styles did not vary between treatment level (strain by stress treatment interaction: *F*_l,36_=0.067, *p* = 0.797, 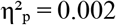). We observed that the stressed fish tended to have higher SOD levels compared to the baseline fish (stress treatment effect: *F*_l,35_ =3.711, *p* = 0.062, 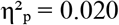) (Fig. 1e). Also, the stressed reactive fish had higher SOD levels compared to the proactive fish (strain effect: *F*_l,35_ =6.881, *p* = 0.013, 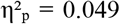). This stress coping style effect did not vary between treatment levels (strain x stress treatment interaction: *F*_l,35_ =0.850, *p* = 0.363, 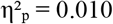).

### Integrated Effect of Acute Stress and Coping Styles on Redox Profiles

The PCA revealed two major axes of variation across all five biomarkers (Fig. 2a). Total GSH, SOD, and ORAC all loaded positively on the first principal component and accounted for 39.99% of the variation (antioxidant axis). GSH ratio and DNA damage loaded positively on the second principal component and accounted for an additional 22.5% of the variation (oxidative stress axis). We found that the stressed fish was tended to have higher PC1 scores compared to the baseline fish (stress treatment effect: *F*_l,36_ = 0.356, *p* = 0.555, 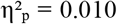). The reactive fish had significantly higher PC1 scores than proactive fish (strain effect: *F*_l,37_ = 14.396, p = 0.001, 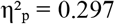) (Fig. 2b). Also, the stressed fish had significantly higher PC2 scores compared with the baseline fish (stress treatment effect: *F*_l,37_ =9.116, p = 0.005, 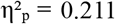) (Fig. 2c). For the behavior variables, stress coping style did not have an effect on time frozen (strain effect: *F*_l,l8_= 0.428, *p* = 0.521, 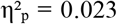), average swimming speed (strain effect: *F*_l,l8_ = 2.685, *p* = 0.119, 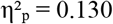), or distance swam (strain effect: *F*_l,l8_ = 2.424, *p* = 0.137, 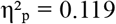). We found a trend for main effects of DNA damage on time frozen only (DNA damage: *F*_l,l5_ = 3.538, *p* = 0.080). There were no significant relationships between levels of other redox biomarker (discrete or composite), stress coping style, or redox biomarker by stress coping style interaction effect on variation of any behavior (Supplementary Table 3).

**Figure 2.**
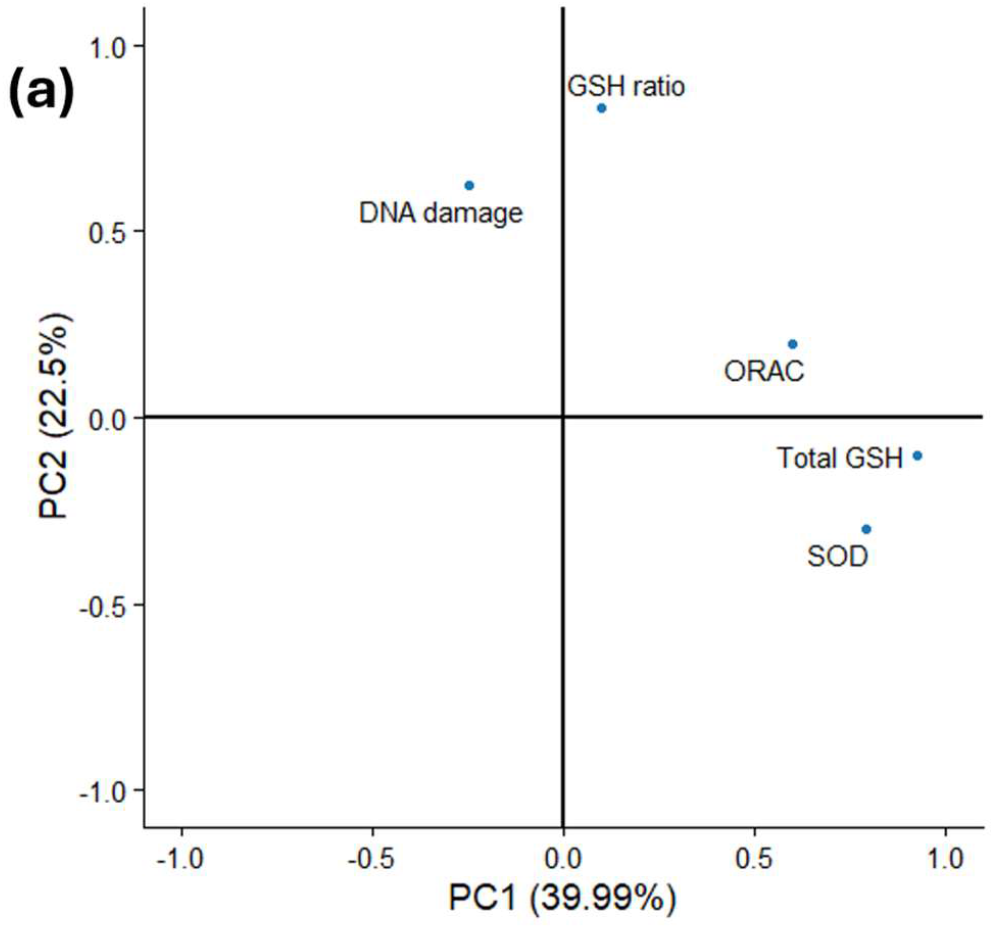
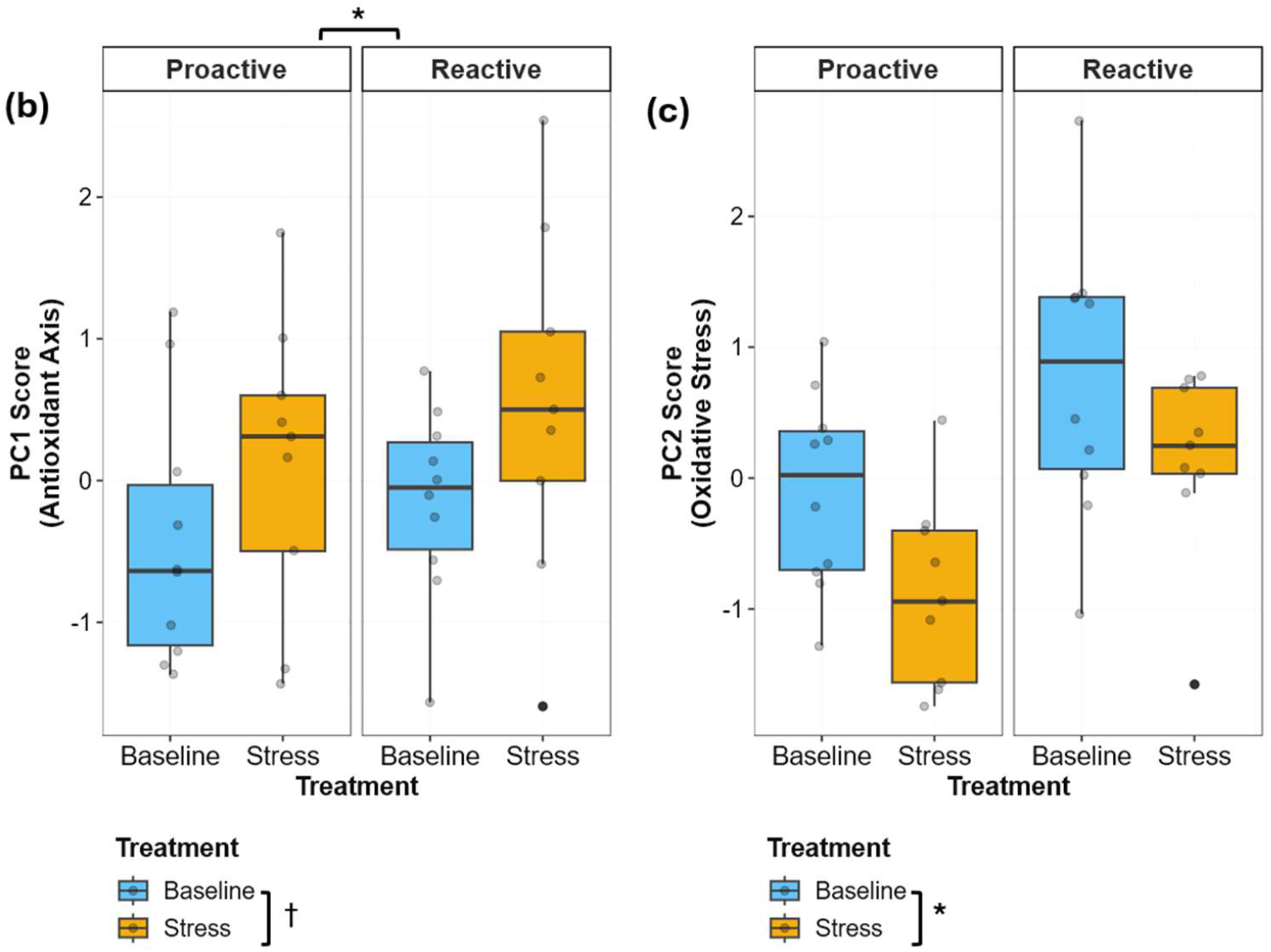
Composite measures (principal component scores) of oxidative stress and antioxidant biomarkers across stress coping styles. (a) Plots of principal components (b) SOD, total glutathione, and ORAC loaded onto principal component 1 (antioxidant axis), whereas (c) DNA damage and glutathione ratio loaded onto principal component 2 (oxidative stress axis). (*) represents *p* < 0.05 and (†) represents 0.05 < *p* < 0.10. Each dot represents a fish.

## Discussion

This study investigated the effects of acute neuroendocrine stress on brain oxidative profiles and how these effects are influenced by individual differences in stress coping style. Our findings show that stress coping style and responses to acute neuroendocrine stressor alters discrete and composite measures of brain oxidative stress and antioxidant markers. More specifically, higher levels of antioxidants (SOD, total glutathione, ORAC, and PC1) in reactive individuals suggest that an individual’s stress-coping style can directly modulate the antioxidant system response to neuroendocrine stressors to potentially mitigate cellular oxidative damage in the brain.

Our study showed that an acute neuroendocrine stressor increases markers of oxidative stress in the brain. Stressed fish tended to have higher amounts of DNA damage and glutathione ratio in both coping styles, suggesting increased molecular damage and altered redox balance in the system via dysregulation of glutathione redox status (Fig. 1). Similarly, the composite measure of oxidative stress (PC2) significantly increased in stressed fish (Fig. 2). Our observations of acute neuroendocrine stress induction of oxidative stress are consistent with other vertebrates, including birds[38] and rodents [39]. Given the evidence that activation of the neuroendocrine stress axis can alter redox balance[7,40,41], we speculate that an excessive release of glucocorticoids disrupted the redox homeostasis in the brain by increasing ROS production, leading to higher glutathione ratios and DNA damage [38].

The onset of oxidative stress is not only indicated by increases in pro-oxidants but can also be a result of decreases in antioxidants [42,43]. We found that across both coping styles, relative to control, there was a significant decrease in antioxidant biomarker levels as well as in the composite measures (PC1 scores) (Fig. 1 & 2). This suggests that neuroendocrine stress can also suppress the antioxidant defense system. Our findings are consistent with other studies showing that acute neuroendocrine stressors in rodents decrease antioxidant levels in both plasma and the brain [40,44,45]. One possible explanation posits that acute stress increases oxidative demand, which can deplete or shift antioxidant pools, thereby reducing measurable antioxidant capacity [44,45].

In addition to overall stress treatment effects on oxidative stress and antioxidant levels, we observed coping-style specific differences. Fish with the reactive stress coping style had significantly higher brain antioxidant activity than the proactive fish across all three biomarkers (total glutathione, total antioxidant capacity, SOD) and the composite measure (PC1 score) (Fig. 1 & 2). Our findings are consistent with prior studies showing higher serum antioxidant capacity in reactive-like mice [17,46,47]. However, opposite patterns have been reported in reactive-like birds, which showed lower plasma antioxidant levels [20]. Notably, while previous studies have largely used peripheral tissues like serum and plasma, our study used whole-brain tissue to measure redox biomarkers. This distinction is important because, relative to other organs, the brain is highly susceptible to oxidative stress due to its high metabolic demand and lipid-rich composition [27,48–50]. Further, the redox profiles between peripheral tissue and brain are not always correlated [24–26]. With the brain’s critical role in modulating behavioral displays and physiological responses, changes in its oxidative stress and antioxidant states may facilitate the biases seen in a stress coping style.

SOD and glutathione are essential for neutralizing ROS, and their levels can be a proxy for antioxidant capabilities [51,52]. ORAC provides an overall measure of antioxidant capacity, including small molecule and enzymatic antioxidants [53,54]. We hypothesize that elevated antioxidant capabilities and capacities in the brain of those with a reactive stress coping style are related to their heightened sensitivity to neuroendocrine stressors. Many studies have shown that the reactive stress coping style have higher baseline, faster release rate, or higher peak cortisol levels than proactive individuals[55–58]. Reactive individuals’ higher antioxidant capacity and capabilities may help mitigate the impact of more frequent cortisol-induced pro-oxidant production in their brains (i.e., oxidative stress). We detected no relationship between individual variation in stress behavior and oxidative stress or antioxidant brain marker levels (Supplementary Table 3). While there may be no linear relationship between stress behaviors and redox biomarkers, we cannot rule out that the lack of differences in neuroendocrine-induced stress behavior between the stress coping styles could be a confound. The inconsistency of the behavioral observations of the current study with prior studies using these selectively bred lines [13,30] could be due to natural variation within the lines.

Overall, this study addresses an important knowledge gap related to whether neuroendocrine stress alters brain oxidative profiles between stress coping styles. The key distinction between coping styles is their antioxidant capability and capacity in response to acute neuroendocrine stressors. Individuals with the reactive stress coping style may have higher brain antioxidant capabilities and capacities to mitigate the presumably more frequent neuroendocrine stress-induced oxidative stress during daily activities. It is unclear how stressor duration may impact this relationship between stress coping styles. Future studies should explore the impacts of chronic neuroendocrine stress on brain redox balance differences between the stress coping styles. Ultimately, our study highlights that the brain’s redox profile may be a key trait that is part of the correlated suite of behaviors and physiological responses to neuroendocrine stressors that characterize a stress coping style.

## Acknowledgements

We thank the UNO ACUP staff for helping with fish husbandry. We are grateful to the Wong lab members for helpful discussion and feedback on earlier versions of this manuscript.

## Animal Ethics and Consent to Participate

Ethical approval for this study was obtained from the University of Nebraska at Omaha’s Institutional Animal Care and Use Committee, IACUC (17-070-09-FC). The present study followed national and institutional guidelines for humane animal treatment and complied with relevant legislation from IACUC.

## Author Contributions

PSJ: Data analysis and interpretation; manuscript drafting.

RYW: Project conceptualization and oversight; data collection; funding acquisition; data interpretation; manuscript drafting.

RJF: Data collection and analysis; manuscript drafting BJB: Data collection and analysis; manuscript drafting

PD: Data analysis and interpretation; funding acquisition; manuscript drafting

