## Supplementary Tables for "Neuroendocrine Stress Induces Differential Oxidative Stress and Antioxidant Profiles between Proactive and Reactive Stress Coping Styles"

**Authors Affiliations:**


### Supplementary Tables

**Table 1:** Two-way ANOVA (Type III sums of squares) testing the effects of sex, stress-coping styles (strain), treatment (baseline vs. stressed), and their interaction on oxidative biomarkers. Values shown are F-tests and Partial Eta Squared ($\eta^2_p$) as effect size.

| **Biomarker** | **Strain Effect** | | | **Sex Effect** | | | **Treatment Effect** | | | **Strain *Treatment*Sex Effect** | | |
|---|---|---|---|---|---|---|---|---|---|---|---|---|
| | **F** | **Sig** | **($\eta^2_p$)** | **F** | **Sig** | **($\eta^2_p$)** | **F** | **Sig** | **($\eta^2_p$)** | **F** | **Sig** | **($\eta^2_p$)** |
| **DNA damage** | 1.264 | 0.27 | 0.039 | 0.039 | 0.845 | 0.001 | 3.83 | 0.059 | 0.11 | 1.418 | 0.251 | 0.155 |
| **Glutathione ratio** | 0.66 | 0.423 | 0.02 | 0.008 | 0.93 | 0 | 2.831 | 0.102 | 0.081 | 0.48 | 0.75 | 0.057 |
| **Total glutathione** | 14.251 | 0.001 | 0.308 | 0.21 | 0.65 | 0.007 | 3.461 | 0.072 | 0.098 | 1.474 | 0.233 | 0.156 |
| **SOD** | 5.717 | 0.023 | 0.156 | 0.111 | 0.742 | 0.004 | 3.125 | 0.087 | 0.092 | 0.896 | 0.478 | 0.104 |

**Table 2:** Two-way ANOVA (Type III sums of squares) testing the effects of sex, stress coping styles (strain), and their interaction with treatment on behavior markers. Values shown are F-tests and Partial Eta Squared ($\eta^2_p$) as effect size.

| Behavior | Strain Effect | | | Sex Effect | | | Strain *Treatment*Sex Effect | | |
|---|---|---|---|---|---|---|---|---|---|
| | F | Sig | ($\eta^2_p$) | F | Sig | ($\eta^2_p$) | F | Sig | ($\eta^2_p$) |
| **Total distance swam (cm)** | 3.825 | 0.068 | 0.193 | 2.145 | 0.162 | 0.118 | 0.118 | 0.736 | 0.007 |
| **Average swim speed (cm/sec)** | 4.055 | 0.061 | 0.202 | 2.019 | 0.175 | 0.112 | 0.111 | 0.744 | 0.007 |
| **Time frozen (secs)** | 1.381 | 0.257 | 0.079 | 2.251 | 0.153 | 0.123 | 0.815 | 0.38 | 0.048 |

**Table 3:** Linear Mixed Model testing the effects of stress coping styles (strain), oxidative damage, antioxidant biomarkers, composite measures, and their interaction on behavior markers. Values shown are F-tests and P-values.

| Behavior | DNA Damage | | | | | | Glutathione Ratio | | | | | | ORAC | | | | | | Total Glutathione | | | | | | SOD | | | | | | PC1 | | | PC2 | | |
|---|---|---|---|---|---|---|---|---|---|---|---|---|---|---|---|---|---|---|---|---|---|---|---|---|---|---|---|---|---|---|---|---|---|---|---|---|
| | Strain | | Biomarker | | Strain *DNA Damage | | Strain | | Biomarker | | Strain*Glutathione Ratio | | Strain | | Biomarker | | Strain *ORAC | | Strain | | Biomarker | | Strain*Total Glutathione | | Strain | | Biomarker | | Strain*SOD | | Strain | Biomarker | Strain*PC1 | Strain | Biomarker | Strain*PC2 |
| | F | Sig | F | Sig | F | Sig | F | Sig | F | Sig | F | Sig | F | Sig | F | Sig | F | Sig | F | Sig | F | Sig | F | Sig | F | Sig | F | Sig | F | Sig | F | Sig | F | F | Sig | F |
| Total distance swam (cm) | 0.374 | 0.55 | 0.629 | 0.44 | 0.248 | 0.625 | 1.013 | 0.329 | 1.118 | 0.306 | 0.087 | 0.772 | 0.999 | 0.332 | 1.834 | 0.195 | 0.819 | 0.379 | 0.125 | 0.728 | 0.583 | 0.456 | 0.125 | 0.728 | 0.006 | 0.938 | 0.068 | 0.798 | 0.097 | 0.76 | 0.536 | 0.476 | 0.058 | 1.952 | 0.184 | 0.468 |
| Average swim speed (cm/sec) | 0.167 | 0.689 | 0.939 | 0.348 | 0.082 | 0.779 | 0.928 | 0.35 | 0.986 | 0.336 | 0.598 | 0.451 | 1.05 | 0.321 | 1.87 | 0.19 | 0.852 | 0.37 | 0.115 | 0.738 | 0.581 | 0.457 | 0.115 | 0.738 | 0.001 | 0.974 | 0.023 | 0.881 | 0.043 | 0.838 | 0.498 | 0.492 | 0.118 | 1.939 | 0.186 | 0.317 |
| Time frozen (secs) | 0.021 | 0.887 | 3.538 | 0.08 | 0.032 | 0.86 | 0.087 | 0.772 | 0.598 | 0.451 | 0.026 | 0.874 | 0.436 | 0.518 | 1.283 | 0.274 | 0.412 | 0.53 | 0.223 | 0.643 | 1.074 | 0.316 | 0.223 | 0.643 | 0.002 | 0.967 | 0 | 0.987 | 0.002 | 0.966 | 0.011 | 0.918 | 0.19 | 0.18 | 0.678 | 0 |

**Table 4: Behavior and Biomarker Data**

| Strain | Sex | Treatment | Total Distance Swam (cm) | Average Swim Speed (cm/secs) | Time Frozen (secs) | DNA Damage (ng 8-OHdG/ng DNA) | GSH Ratio (reduced: oxidized) | ORAC (umol TE/Ug protein) | SOD (U/mg protein) | Total GSH (Total glutathione /Ug protein) | PC1 | PC2 |
|---|---|---|---|---|---|---|---|---|---|---|---|---|
| Reactive | Male | Baseline | - | - | - | 0.087 | 4.0398 | 2.2698 | 0.9279 | 6.14 | 0.70428 | -0.41866 |
| Reactive | Female | Baseline | - | - | - | 0.0872 | 3.4058 | 2.7054 | 0.878 | 5.75 | 0.91341 | -0.48179 |
| Reactive | Male | Baseline | - | - | - | 0.1105 | 3.3889 | 2.2623 | 0.8673 | 4.42 | -0.38983 | -0.29219 |
| Reactive | Female | Baseline | - | - | - | 0.1047 | 3.7962 | 2.5262 | 0.7111 | 6.1 | 0.459 | 0.27544 |
| Reactive | Female | Baseline | - | - | - | 0.11 | 2.5291 | 3.093 | 1.0885 | 6.37 | 1.99416 | -0.44227 |
| Reactive | Male | Baseline | - | - | - | 0.1182 | 2.2887 | 2.3053 | 0.6053 | 4.67 | -1.06196 | -0.73595 |
| Reactive | Female | Baseline | - | - | - | 0.099 | 4.48 | 2.5342 | 1.0959 | 8.71 | 2.6639 | 0.34957 |
| Reactive | Male | Baseline | - | - | - | 0.12 | 2.893 | 2.5536 | 1.1765 | 7.11 | 1.807 | -0.46926 |
| Reactive | Male | Stressed | 102.603 | 0.3468 | 286.987 | 0.0838 | 4.7138 | 2.6124 | 0.7639 | 5.81 | 0.8068 | 0.53073 |
| Reactive | Male | Stressed | 1384.34 | 4.6801 | 13.1798 | 0.1642 | 5.1569 | 2.9565 | 0.7133 | 4.77 | 0.61365 | 3.36916 |
| Reactive | Male | Stressed | 24.6745 | 0.0834 | 294.027 | 0.0859 | 3.3707 | 2.4827 | 0.6737 | 5.5 | 0.00835 | -0.57345 |
| Reactive | Female | Stressed | 2.2961 | 0.0078 | 299.967 | 0.1184 | 4.8519 | 2.7492 | 0.6329 | 4.55 | 0.0848 | 1.79872 |
| Reactive | Male | Stressed | 1444.6 | 4.9361 | 0.0334 | 0.1169 | 2.055 | 2.1097 | 0.5664 | 3.57 | -1.95809 | -1.10939 |
| Reactive | Male | Stressed | 808.654 | 2.7332 | 87.9877 | 0.1298 | 3.2295 | 2.4048 | 1.0355 | 6.41 | 1.00161 | 0.05233 |
| Reactive | Female | Stressed | 26.8386 | 0.0908 | 294.962 | 0.0822 | 3.403 | 2.7569 | 0.5836 | 5.54 | 0.22102 | -0.30293 |
| Reactive | Male | Stressed | 26.8545 | 0.0908 | 295.028 | - | 2.7378 | 2.0847 | 1.0247 | 6.28 | - | - |
| Reactive | Female | Stressed | 4.0246 | 0.0136 | 298.564 | 0.0971 | 3.9123 | 2.4972 | 0.9043 | 5.28 | 0.54534 | -0.02024 |
| Reactive | Male | Stressed | 1661.95 | 5.6085 | 43.2097 | 0.1128 | 3.7084 | 2.3017 | 0.8762 | 5.71 | 0.31381 | 0.05647 |
| Reactive | Male | Baseline | - | - | - | 0.0927 | 3.1571 | 2.2217 | 0.905 | 5.24 | 0.00018 | -1.02537 |
| Reactive | Female | Baseline | - | - | - | 0.0997 | 3.2162 | 2.6858 | 0.9425 | 6.35 | 1.228 | -0.3924 |
| Proactive | Female | Baseline | - | - | - | 0.0832 | 2.3882 | 2.5538 | 0.7527 | 5.64 | 0.16508 | -1.46404 |
| Proactive | Female | Baseline | - | - | - | 0.1073 | 3.0519 | 2.4699 | 0.8635 | 6.11 | 0.58506 | -0.46457 |
| Proactive | Female | Baseline | - | - | - | 0.1276 | 2.7371 | 2.504 | 0.9328 | 5.92 | 0.595 | -0.22776 |
| Proactive | Male | Baseline | - | - | - | 0.0791 | 2.6244 | 2.665 | 0.7411 | 5.2 | 0.1601 | -1.25871 |
| Proactive | Male | Baseline | - | - | - | 0.0949 | 5.1212 | 2.2578 | 0.4799 | 4.05 | -1.07469 | 1.08524 |
| Proactive | Female | Baseline | - | - | - | 0.1461 | 3.5628 | 2.541 | 0.6373 | 4.37 | -0.60117 | 1.24698 |
| Proactive | Male | Baseline | - | - | - | 0.099 | 2.2559 | 2.4024 | 0.8126 | 3.89 | -0.74913 | -1.35376 |
| Proactive | Female | Baseline | - | - | - | 0.1085 | 2.2167 | 2.5039 | 0.5975 | 4.59 | -0.82025 | -0.85381 |
| Proactive | Female | Baseline | - | - | - | 0.1184 | 3.0569 | 2.0074 | 0.9197 | 4.29 | -0.77562 | -0.65126 |
| Proactive | Female | Baseline | - | - | - | 0.1048 | 2.5298 | 2.2375 | 0.6892 | 3.72 | -1.30463 | -1.01777 |
| Proactive | Male | Stressed | 2347.81 | 8.0096 | 0.0667 | 0.12 | 4.0639 | 2.0276 | 0.8277 | 5.7 | -0.14009 | 0.32562 |
| Proactive | Female | Stressed | 2415.1 | 8.3235 | 0 | 0.1171 | 3.1434 | 2.4725 | 0.7043 | 4.76 | -0.3829 | 0.01658 |
| Proactive | Female | Stressed | 1966.94 | 6.6707 | 82.1819 | 0.116 | 1.7864 | 2.274 | - | 2.06 | - | - |
| Proactive | Female | Stressed | 11.7098 | 0.0396 | 299.967 | 0.1355 | 3.5708 | 2.5299 | 0.6341 | 3.56 | -0.95552 | 0.97545 |
| Proactive | Female | Stressed | 1266.27 | 5.1845 | 30.9308 | 0.1507 | 4.0452 | 2.3461 | 0.5934 | 4.29 | -0.92924 | 1.62445 |
| Proactive | Male | Stressed | 2590.43 | 9.4286 | 0.367 | 0.1238 | 3.6679 | 2.0847 | 0.6142 | 4.1 | -1.34564 | 0.34594 |
| Proactive | Female | Stressed | 2.5001 | 0.0084 | 299.967 | 0.1131 | 5.0167 | 2.2435 | 0.4824 | 3.73 | -1.29525 | 1.45947 |
| Proactive | Male | Stressed | 739.344 | 2.4984 | 230.997 | 0.118 | 2.4211 | 2.3939 | 0.7479 | 4.21 | -0.78303 | -0.67177 |
| Proactive | Female | Stressed | 263.74 | 0.8922 | 285.152 | 0.109 | 3.4044 | 2.5732 | 0.805 | 4.83 | 0.09581 | 0.02812 |
| Proactive | Male | Stressed | 178.284 | 0.6019 | 282.749 | 0.1167 | 4.5144 | 2.0952 | 0.8227 | 4.7 | -0.39932 | 0.68709 |
